# Silencing long ascending propriospinal neurons after spinal cord injury improves hindlimb stepping in the adult rat

**DOI:** 10.1101/2021.05.18.444653

**Authors:** Courtney T. Shepard, Amanda M. Pocratsky, Brandon L. Brown, Morgan A. Van Rijswijck, Rachel M. Zalla, Darlene A. Burke, Johnny M. Morehouse, Amberly S. Reigler, Scott R. Whittemore, David S.K. Magnuson

**Affiliations:** Interdisciplinary Program in Translational Neuroscience, School of Interdisciplinary and Graduate Studies, University of Louisville, Louisville, Kentucky 40292, USA; Dept. of Anatomical Sciences and Neurobiology, University of Louisville, Louisville, Kentucky 40292, USA; Kentucky Spinal Cord Injury Research Center, University of Louisville, Louisville, Kentucky 40292, USA; Speed School of Engineering, University of Louisville, Louisville, Kentucky 40292, USA; Dept. of Neurological Surgery, University of Louisville, Louisville, Kentucky 40292, USA

**Keywords:** spinal cord injury, propriospinal neurons, locomotion, viral vector, neuronal silencing

## Abstract

Long ascending propriospinal neurons (LAPNs) are a subpopulation of spinal cord interneurons that directly connect the lumbar and cervical enlargements. In uninjured animals, conditionally silencing LAPNs resulted in disrupted left-right coordination of the hindlimbs and forelimbs in a context-dependent manner, demonstrating that LAPNs secure alternation of the fore- and hindlimb pairs during overground stepping in the adult rat. Given their ventrolateral location in the spinal cord white matter, many LAPN axons likely remain intact following thoracic spinal cord injury (SCI), suggesting a potential role in the recovery of stepping. Thus, we hypothesized that silencing LAPNs after SCI would result in diminished hindlimb locomotor function. We found instead that silencing of spared LAPNs post-SCI restored the left-right hindlimb coordination associated with alternating gaits that was lost as a result of SCI. Several other fundamental characteristics of hindlimb stepping were also improved and the number of abnormal steps were reduced. However, hindlimb-forelimb coordination was not restored. These data suggest that the temporal information carried between the enlargements by the LAPNs after SCI may be detrimental to hindlimb locomotor function. These observations have implications for our understanding of the relationship between injury severity and functional outcome, for the efforts to develop neuro- and axo-protective therapeutic strategies, and also for the clinical study/implementation of spinal stimulation and neuromodulation.

## Introduction

In mammals, locomotion involves descending commands from supraspinal centers and peripheral input from sensory systems converging on spinal locomotor circuitry. This circuitry includes central pattern generators (CPGs), intrinsic spinal networks that generate the coordinated muscle activity associated with stereotypic limb movements during stepping *(Sherrington and Laslett, 1903; Sherrington, 1910, Graham Brown, 1911)*. Each limb has its own CPG and the lumbar and cervical CPG pairs are interconnected by long-ascending (LAPNs) and long-descending (LDPNs) propriospinal neurons *(Giovanelli Barilari and Kuypers 1969; English et al., 1985)*. These neurons provide the functional coupling of the two enlargements allowing precise temporal information to be passed between the hindlimb and forelimb CPGs *(Miller et a., 1975; English, 1979; Rossignol et al., 1993; Juvin et al., 2005, 2012)*. LAPN somata reside in the intermediate gray matter, primarily in laminae VII and VIII, with 40-60% having commissural axons *(Reed et al., 2006, 2009; Brockett et al., 2013)* that cross at-level before ascending in the outermost regions of the lateral and ventrolateral funiculi *(Reed et al., 2006, 2009; Basso et al., 2002)*.

Spinal cord injury (SCI) disrupts communication between the brain and spinal cord, resulting in an immediate inability to initiate and maintain patterned weight-supported locomotion at or below the level of lesion *(Dietz and Harkema, 2004; Fong et al., 2009; Côté et al., 2017)*. Even if classified as neurologically complete, most SCIs are anatomically incomplete with some sparing of white matter, most often the outermost rim of the lateral and ventrolateral funiculi where LAPN axons reside *(Brockett et al, 2013)*. These neurons and their axons comprise a percentage of the anatomically spared circuitry, thus providing a functional bridge across the injury site, making them well-suited to participate in locomotor recovery after incomplete thoracic SCI *(Conta and Stelzner, 2004; Conta et al., 2010, 2011; Siebert et al., 2010)*. Many studies have reinforced this notion, suggesting that propriospinals, albeit descending in these studies, can contribute to locomotor recovery by serving as injury-crossing bridges *(Bareyre et al., 2004; Vavrek et al., 2006; Flynn et al., 2011; Filli et al., 2014; Benthall et al., 2017)*.

Conditionally silencing LAPNs in the uninjured rat resulted in the partial decoupling of the forelimb and hindlimb pairs, disrupting alternation at each girdle *(Pocratsky et al., 2020)*. These changes were independent of locomotor rhythm and speed and did not affect intralimb joint movements or coordination. Given their anatomical characteristics and functional importance, we hypothesized that LAPNs contribute to functional recovery after incomplete thoracic SCI and that silencing them would further disrupt locomotion by reducing the number of functional spared axons that cross the injury site leaving animals unable to produce a functional stepping pattern.

## Results

### Silencing alters interlimb coordination while other key features of locomotion are maintained

We marked the skin overlying the iliac crest, hip, ankle, and toe (Figure 1 – figure supplement 1a,c) to assess intralimb coordination of the hip/knee (proximal) and knee/ankle (distal) segment angles. In uninjured animals, we observed normal rhythmic excursions of the proximal and distal limb segments (Figure 1 – figure supplement 1b,d,h), as well as coordinated flexor-extensor movements of the proximal and distal angles during normal walking (Figure 1 – figure supplement 1e-g).

We examined the coupling patterns of various limb pairs by dividing the initial contact time of one limb by the stride time of the second limb, expressed as a phase value. Phase values of 0 or 1 indicate synchrony (lead-limb dependent) and values of 0.5 indicate alternation, typically shown on circular plots as 0 or 1 at 0° and 0.5 at 180°. For slower gaits such as walk and trot, phase values of hindlimb, forelimb, and homolateral hindlimb-forelimb pairs would be concentrated around 0.5, indicating alternation, while the heterolateral hindlimb-forelimb pair would concentrate around 0 or 1, indicating synchrony (Fig 1c_i_). These values are reversed for bounding behavior, with synchrony of the hindlimb and forelimb pairs, and alternation of the homolateral and heterolateral limb pairs (Fig 1c_ii_). To eliminate any discrepancies between lead limb selection in the animals, phase values were converted from a circular scale (0-1) to a linear scale (0.5-1) (Fig 1d).

**Figure 1.**
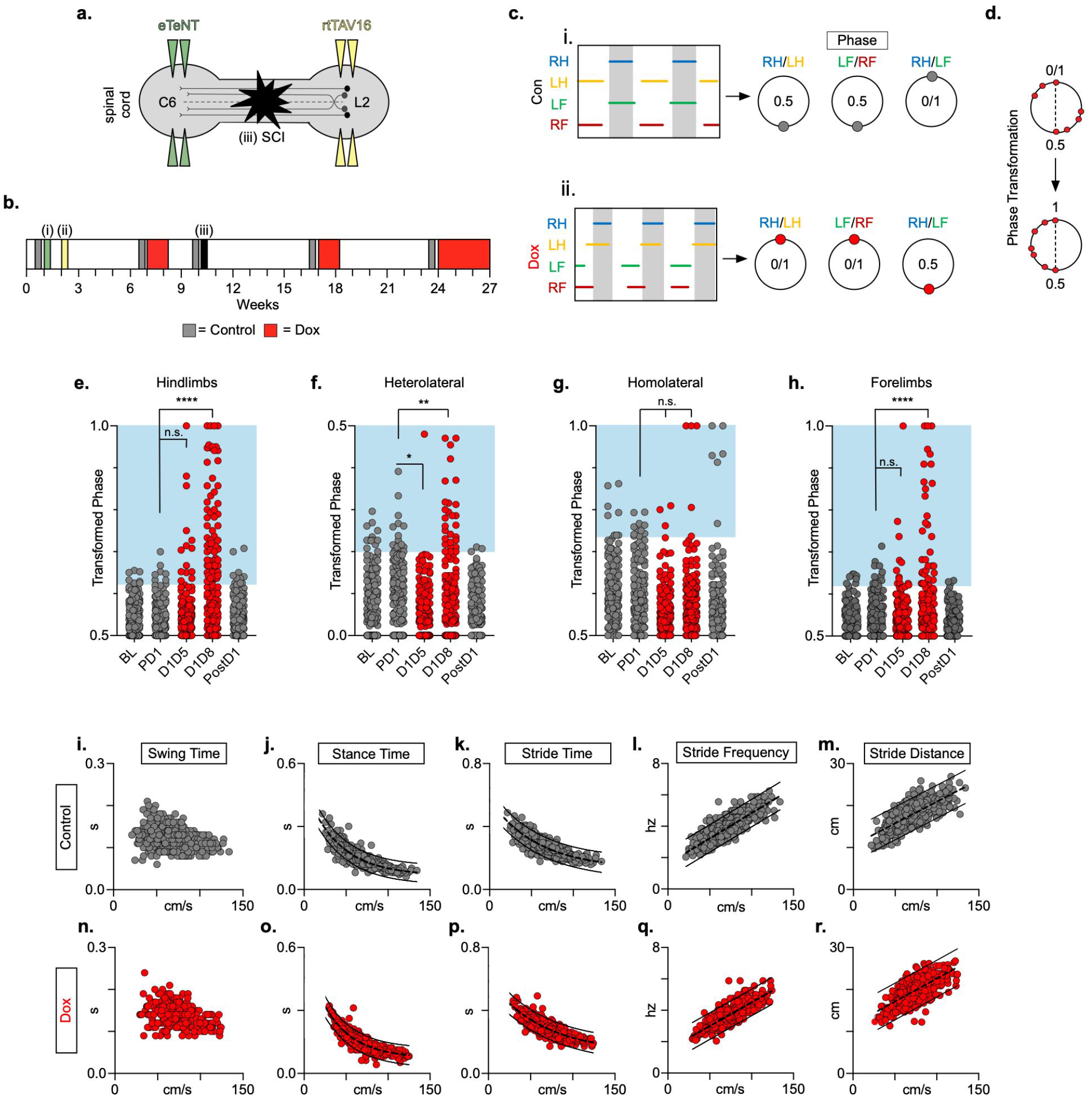
Silencing LAPNs disrupts interlimb coordination without affecting key features of locomotion. (**a**) Bilateral injections of enhanced tetanus neurotoxin (eTeNT, green) and Tet-On rtTAV16 (yellow) were performed at C6 and L2 spinal cord levels, respectively, followed by a SCI (black star). Administration of doxycycline (Dox, red boxes) induces eTeNT expression in doubly-infected neurons. eTeNT is transported to the cell terminals where it prevents synaptic vesicle release into the synaptic cleft, effectively silencing neurotransmission in the targeted neuronal *(Pocratsky et al., 2017a; Kinoshita et al., 2012)*. Following viral injections (i) and (ii), pre-injury behavioral assessments were taken at three control time points and a single round of Dox administration. Following SCI (iii, black star), behavioral assessments were repeated at two control time points and during two rounds of Dox administration (grey boxes) (**b**). Representative foot fall graphs are shown with corresponding coordination phase values for both Control and Dox timepoints (**c;** RH = Right Hindlimb, LH = Left Hindlimb, LF = Left Forelimb, RF = Right Forelimb). Shaded regions indicate swing phase of LH and RF within the footfall cycle. Phase values generated from foot fall graphs were converted to a scale of 0-1 to 0.5-1 for the hindlimb, forelimb, and homolateral hindlimb-forelimb limb pairs and 0-0.5 for the heterolateral hindlimb-forelimb pair so they could be further examined on a linear scale (**d**). The hindlimb and forelimb limb pairs were significantly altered during LAPN silencing, while the heterolateral and homolateral hindlimb-forelimb limb pairs remained unaffected (**e-h**, # steps beyond control variability: PD1 hindlimbs n=7/168 [4.17%] vs D1D5 hindlimbs *n*=14/166 [8.43%]; n.s., *z*=1.62; PD1 hindlimbs *n*=7/168 [4.17%] vs D1D8 hindlimbs *n*=72/161 [44.72%]; *p*<.001, *z*=9.63; PD1 heterolateral *n*=8/168 [4.76%] vs D1D5 heterolateral *n*=1/166 [0.60%]; *p*<.05, *z*=2.38; PD1 heterolateral *n*=8/168 [4.76%] vs D1D8 heterolateral *n*=21/161 [13.04%]; *p*<.01, *z*=2.65; PD1 homolateral *n*=7/168 [4.17%] vs D1D5 homolateral *n*=3/161 [1.80%]; n.s., *z*=1.27; PD1 homolateral *n*=7/168 [4.17%] vs D1D8 homolateral *n*=4/161 [2.48%]; n.s, *z*=0.85; PD1 forelimbs n=14/168 [8.33%] vs D1D5 forelimbs n=12/166 [7.23%]; n.s., *z*=0.38; PD1 forelimbs n=14/168 [8.33%] vs D1D8 forelimbs n=35/161 [21.74%]; *p*=.001, *z*=3.45; Binomial Proportion Test; circles=individual step cycles; shaded region=values beyond control variability). Spatiotemporal measures (swing time, stance time, stride time, stride distance) were plotted against speed for Control (**i-m**) and Dox (**n-r**) timepoints. An exponential decay line of best fit is displayed for stance time and stride time graphs (stance time: Control R^2^=.785 vs Dox R^2^=.735; stride time: Control R^2^=.708 vs Dox R^2^=.667), while a linear line of best fit is displayed for stride distance (stride distance: Control R^2^=.584 vs Dox R^2^=.513; line of best fit indicated by dotted line). 95% prediction intervals are also shown as solid lines.

At control timepoints, hindlimb and forelimb pairs maintained a phase value around 0.5. As expected, phase values for the heterolateral and homolateral hindlimb-forelimb pairs focused around synchrony and alternation, respectively, although with greater variability (Fig 1e-h, grey). We determined the mean phase value of the limb pairs and any value >2 S.D. from this mean was considered “irregular”, as is indicated by the blue boxes. Silencing LAPNs led to disruptions in the left-right alternation of the hindlimb and forelimb pairs, with modest changes to the heterolateral hindlimb-forelimb pair (Fig 1e,f,h, red). The homolateral limb pair remained unaffected (Fig 1g, red). The effects of silencing on limb pair relationships were reversed when Dox was removed from the drinking water (Fig 1e-h, “PostD1”). However, silencing did not affect the well-described, speed-dependent gait indices of swing time (Fig 1i,n), stance time (Fig 1j,o), stride time (Fig 1k,p), stride frequency (Fig 1l,q), or stride distance (Fig 1m,r). The excursion of the proximal limb angle remained unaffected by silencing, whereas there were slight, but significant effects on the excursion of the distal limb angle (Figure 1 – figure supplement 1h-j) *(Pocratsky et al., 2020)*. These findings replicate our earlier observations and show that LAPNs secure interlimb coordination with little to no change in intralimb coordination or other fundamental gait characteristics in an otherwise intact rat.

### Spared LAPNs express synapse-silencing eTeNT.EGFP chronically

The majority of LAPN axons reside within the ventrolateral funiculus (VLF) *(English et al., 1985; Reed et al., 2006; Molenaar and Kuypers, 1978)*. Importantly, we found the average spared white matter at the injury epicenter to be approximately 19% (Fig 2A) and that the outermost rim of the VLF was largely spared in this injury model (Fig 2b-k).

**Figure 2.**
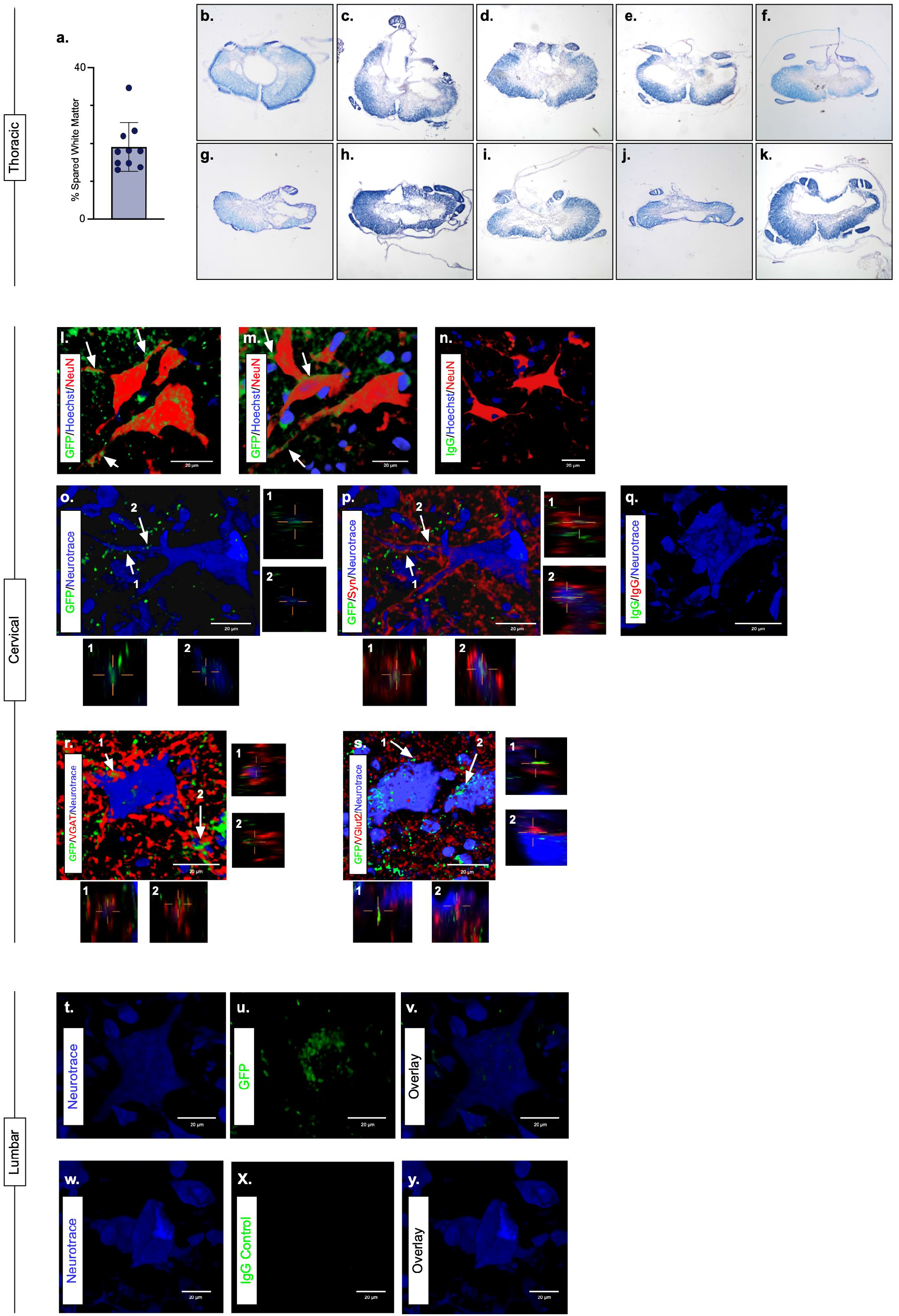
Presence of eTeNT-EGFP in putatively silenced LAPNs across the level of injury. In the thoracic spinal cord, the percentage of spared white matter at the epicenter ranges from 13.01% – 34.64% (**a**). White matter damage at the spinal cord injury epicenter as confirmed by histology (**b-k**). Individual images represent the injury epicenter of each animal used in the main data set (*N*=10; average white matter percentage: 19.02%, S.D. 6.44%). High magnification, volume-rendered images demonstrating eTeNT.EGFP putatively positive fibers (green) surrounding NeuN stained neurons (red) with Hoechst nuclear counterstain (blue) in cervical spinal cord segments of interest (**l,m**) (100x magnification, C6-C7 spinal cord). White arrows indicate areas of colocalization. Isotype control reveals minimal immunoreactivity (**n**, IgG controls for eTeNT.EGFP shown). eTeNT.EGFP (green) signal co-localizes with neuronal processes (blue) and synaptophysin (red) (**o,p**). Isotype controls further show minimal reactivity (**q**, IgG controls of synaptophysin and eTeNT.EGFP shown). eTeNT.EGFP also colocalizes with the inhibitory neurotransmitter marker vesicular GABA transporter (**r**, VGAT, red) and the excitatory neurotransmitter vesicular glutamate transporter 2 (**s**, VGlut2, red). eTeNT.EGFP putative cell bodies (green) in the lumbar spinal cord colocalized with NeuroTrace fluorescent Nissl stained neurons (blue) (**t-v**, L1-L2 spinal cord). Minimal presence of eTeNT.EGFP signal in isotype controls (**w-y**, L1-L2 spinal cord).

Immunohistochemistry to enhance eTeNT.EGFP revealed EGFP-positive fibers (putative axon terminals) surrounding neuronal processes in the caudal cervical enlargement (Fig 2l,m). Similar to uninjured histology (Pocratsky et al 2020), eTeNT.EGFP co-localized with synaptophysin (Fig 2o,p), vesicular GABA transporter (Fig 2r, VGAT), and vesicular glutamate transporter 2 (Fig 2s, VGlut2), markers of synapses and excitatory and inhibitory neurotransmitters, respectively. Isotype controls revealed minimal-to-no immunoreactivity (Fig 2n,q).

In rostral lumbar spinal cord segments, eTeNT-EGFP-positive LAPN cell bodies colocalized with fluorescent Nissl-stained (NeuroTrace) neurons in the intermediate gray matter (Fig 2t-v) *(Reed et al., 2006; Pocratsky et al., 2020)*. Isotype controls showed no immunoreactivity (Fig 2w-y). Taken together, these data suggest that some proportion of LAPNs were spared after moderate T10 contusion injury and that double-infected LAPNs maintained both cell body and axon terminal expression of eTeNT following SCI. Thus, any behavioral changes seen during Dox administration would be concomitant with active eTeNT.EGFP expression in these spared LAPNs.

### Silencing LAPNs post-SCI improves locomotor function

We next explored the effects of LAPN silencing on locomotion after injury. We hypothesized that LAPNs spared after SCI contribute to spontaneous functional recovery, and that silencing them effectively makes the injury more severe and should reduce recovered stepping capacity. As shown in Fig 3a,b, large shifts between the defined alternate and cruciate gait patterns occur post-SCI, but revert to a consistent alternate pattern when LAPNs are silenced. RI and CPI were modestly improved, (Fig 3c,d) as was PSI, signifying a general improvement in paw placement order and timing (Fig 3e). It is important to consider that RI and CPI account for the step sequence pattern of all 4 limbs, while PSI relies on the plantar stepping ratio of the hindlimbs to the forelimbs.

**Figure 3.**
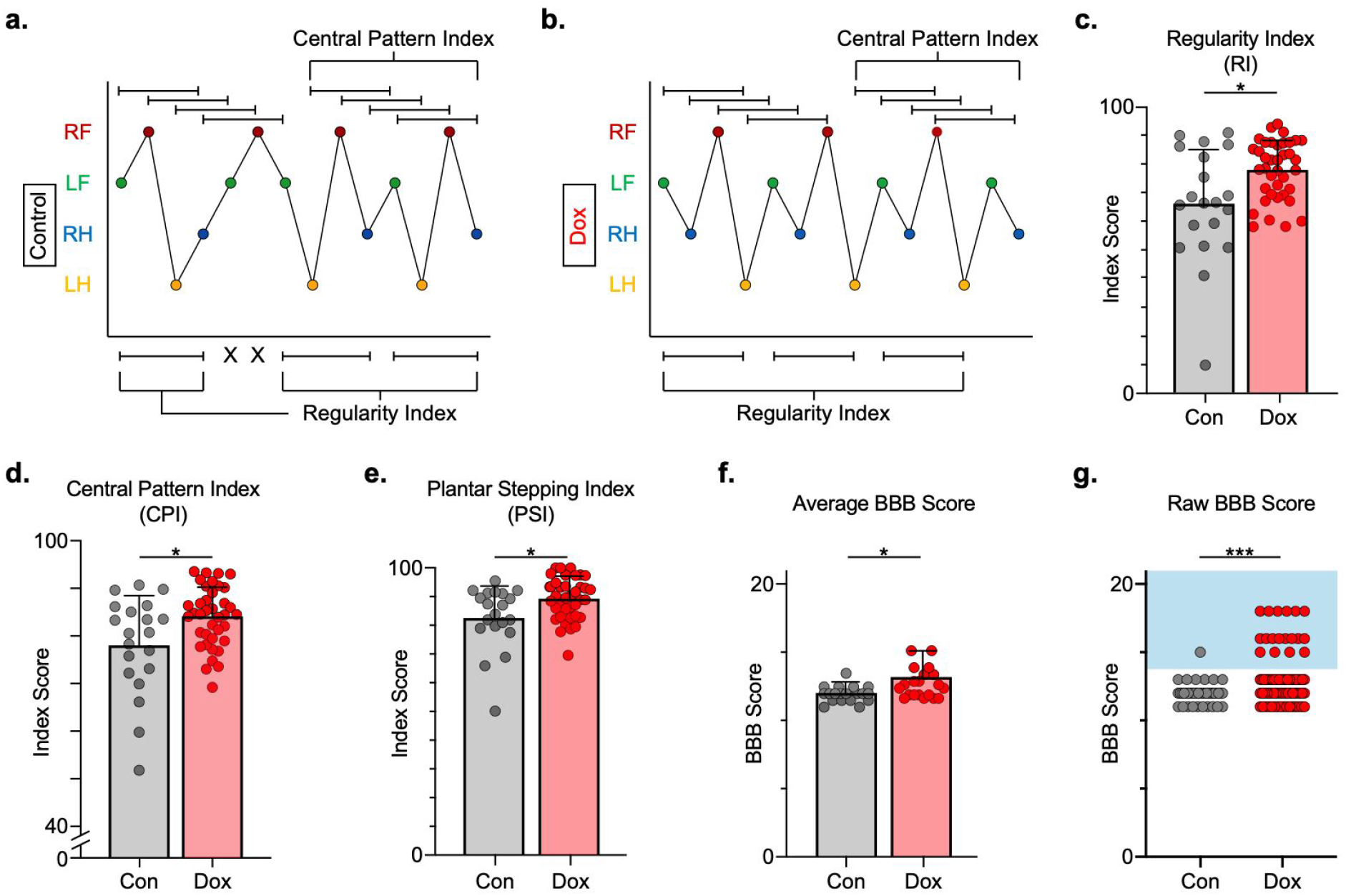
Silencing LAPNs post-SCI restores coordination indices and improves gross locomotor outcomes. Footfall patterns are shown from example stepping passes to demonstrate how gait indices such as regularity index (RI) and central pattern index (CPI) differed between post-SCI Control (**a**) and post-SCI Dox (**b**) stepping (RF = Right Forelimb, LF = Left Forelimb, RH = Right Hindlimb, LH = Left Hindlimb). “X” indicates incorrect footfall order as defined by the typical foot patterns in RI. The average RI scores (**c**, Control RI: 65.74±18.52 vs Dox RI: 76.97±7.70, *t*=2.88, *df*=9, *p*=.018; paired t-test), CPI scores (**d**, Control CPI: 78.37±8.39 vs Dox CPI: 84.84±3.39, *t*=2.94, *df*=9, *p*=.016; paired t-test), and PSI scores (**e**, Control PSI: 82.72±9.97 vs Dox PSI: 89.07±4.87, *t*=2.76, *df*=9, *p*=.022; paired t-test) are demonstrated with individual animal averages (grey circles and red circles for Control and Dox, respectively). Average BBB scores for Control and Dox timepoints are shown (**f**, group average ± S.D. [Control to Dox]; *p*=.663, mixed model ANOVA, Bonferroni *post hoc*). No significant difference was found between right and left BBB scores so they were combined for average and raw score ([Left vs Right]; *p*=.001, mixed model ANOVA, Bonferroni *post hoc*; data not shown). To demonstrate BBB scores prior to averaging, right and left hindlimb raw BBB scores are shown in (**g**, Control: *n*=1/40 [0.025%] vs Dox: *n*=18/76 [23.68%]; *p*<.001, *z*=4.23; Binomial Proportion Test; circles=individual left or right BBB scores; blue shaded region = values beyond post-SCI control variability).

Consistent with previous studies, BBB scores plateaued mid-scale, 11-13 out of 21 *(Basso et al., 2002; Smith et al., 2006)* and increased modestly during silencing (Fig 3f), with a greater proportion of scores for each hindlimb exceeding 13 (range 13-18; Fig 3g). These scores suggest that weight support by the hindlimbs, interlimb coordination and toe clearance are improved during silencing.

### LAPN silencing leads to modest improvements in intralimb coordination after SCI

To better assess the general improvement in locomotor performance suggested above, we next examined hindlimb kinematics during overground stepping post-SCI and following LAPN silencing. After SCI, compromised muscle activation leads to dorsal stepping where animals bear weight on the dorsal surface of their toes or hindpaws (Fig 4a). Proximal and distal limb angle excursions were determined for plantar and dorsal steps throughout each locomotor bout (Fig 4a). Angle excursions were reduced post-SCI during most locomotor bouts, and intralimb coordination was disrupted such that peaks (maximal extension) and troughs (maximal flexion) were no longer occurring simultaneously (Fig 4b,d,e; Video 1). Conditional silencing of spared LAPNs improved the coordination of the proximal and distal hindlimb angles, such that the cyclic properties of each angle were restored (Fig 4c,g,h; Video 2). The excursions of both distal and proximal hindlimb joint angles were also improved (Fig 4f,i), suggesting that the primary effect of silencing may not be on intralimb coordination. These results suggest that changes seen during LAPN silencing after SCI involve modest improvements in the coordination and execution of the hindlimb joint movements.

**Figure 4.**
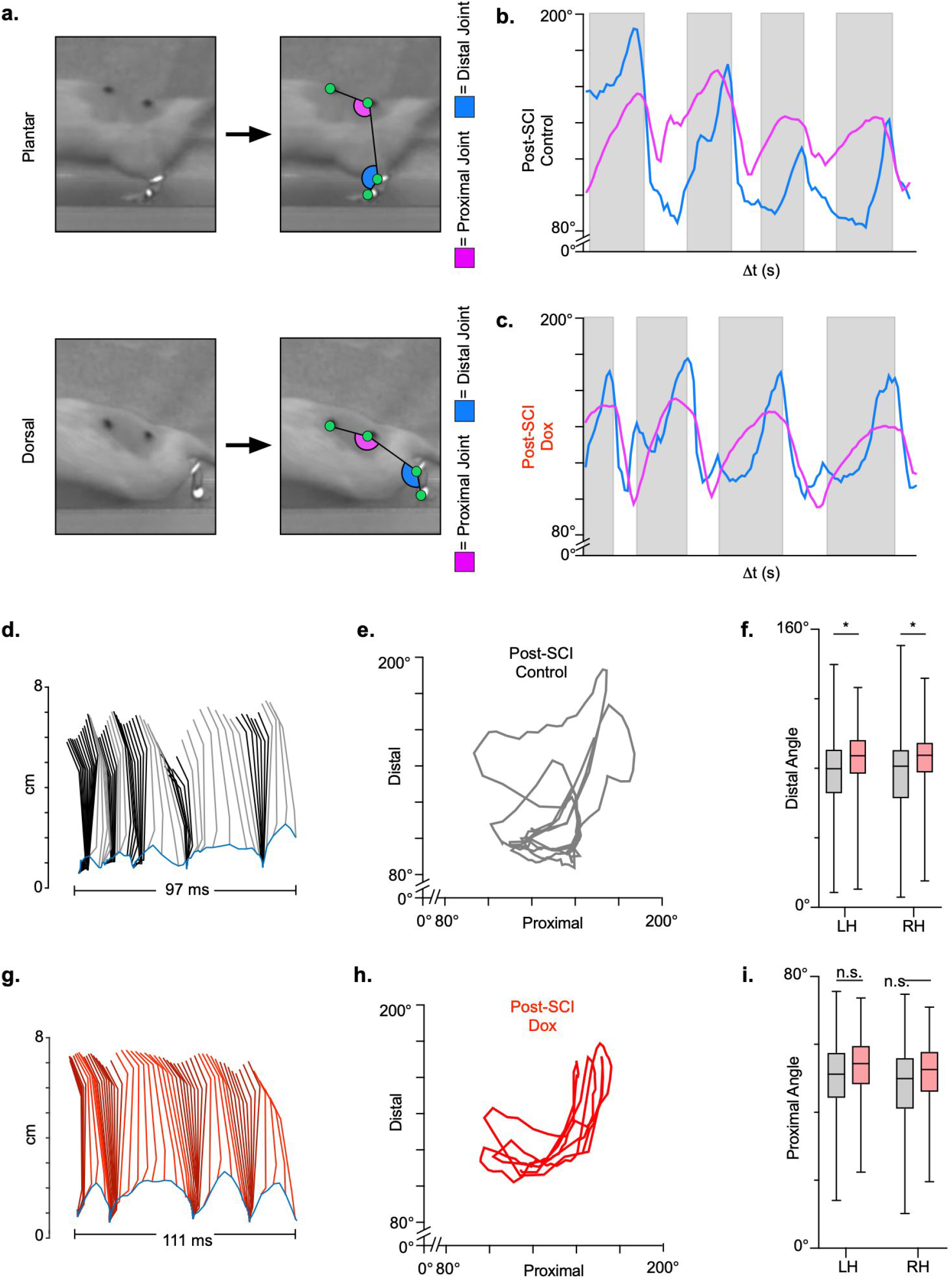
LAPN silencing leads to modest improvements in intralimb coordination after SCI. An example sagittal view of kinematic markers for plantar steps and dorsal steps is shown in (**a**). Excursion traces of the proximal (purple) and distal (blue) joint angles during post-SCI Control (**b**) and Dox (**c**) timepoints are shown. Two-dimensional stick figures of hindlimb stepping (**d,g**) and angle-angle plots (**e,h**) from the same passes shown in (**b,c**). Toe height throughout the pass is indicated as a blue trace on the bottom of the two-dimensional stick figures in **d** and **g**, with the angles around 2cm indicating the distal joint movements (ankle) and the angles around 6cm indicating the proximal joint movements (hip). Grey boxes indicate periods in which the paw is in contact with the surface (stance phase). Angle-angle plots are shown to directly compare the distal joint angle to the proximal joint angle for the same recording frame. In **f,i**, grey boxes indicate post-SCI Control data while pink boxes indicate post-SCI Dox data. Minor improvements in distal joint angle excursion (**f**, Control left: 74.62±11.96 vs Dox left: 85.74±7.24, *t*=3.35, *df*=9, *p*=.009; Control right: 75.29±8.73 vs Dox right: 83.40±6.99, *t*=2.70, *df*=9, *p*=.024; paired t-tests; LH= Left Hindlimb, RH = Right Hindlimb) and proximal joint angle excursion (**i**, Control left: 47.64±7.33 vs Dox left: 51.47±4.33, *t*=2.65, *df*=9, *p*=.027; Control right: 46.45±5.58 vs Dox right: 49.37±3.80, *t*=2.93, *df*=9, *p*=.017; paired t-tests; middle bar indicates group average with extension bars indicating range of raw data) are demonstrated as a result of silencing.

### Hindlimb coupling and paw placement are improved during post-SCI silencing

We quantified dorsal steps in the ventral view based on the appearance of one or more toes, or the paw itself, being curled under to make contact with the walking surface (Fig 5a) *(Keller et al., 2017)*. We then compared the dorsal stepping index (DSI), the ratio of dorsal steps to total hindlimbs steps, at post-SCI control and LAPN silenced timepoints. The DSI was drastically reduced during silencing (11.21±7.97) as compared to control timepoints (24.41±16.1) (Fig 5b). The sidedness of dorsal steps did not change between control and silencing, with ~60% of dorsal steps occurring in the right hindlimb and ~40% of dorsal steps occurring in the left hindlimb during both conditions (Fig 5c). Thus, improvements seen during LAPN silencing were not biased to one side. For the following analyses, we used the left hindlimb as the reference limb.

**Figure 5.**
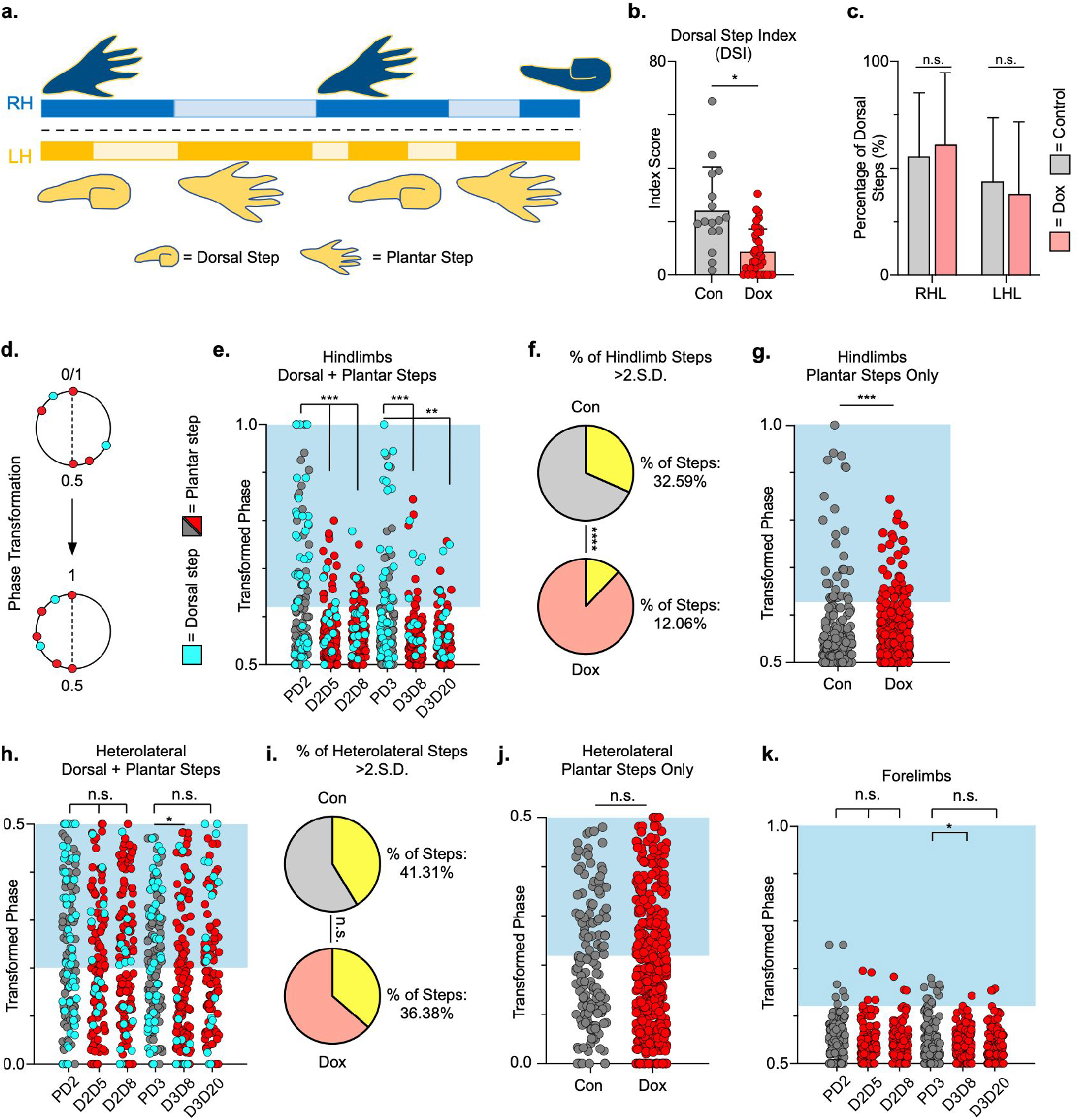
Hindlimb, but not hindlimb-forelimb, coupling relationships are restored during post-SCI silencing. An example pass with multiple dorsal and plantar steps shown in (**a**; LH = Left Hindlimb, light yellow boxes = LH swing phase, dark yellow boxes = LH stance phase; RH = Right Hindlimb, light blue boxes = RH swing phase, dark blue boxes = RH stance phase). Dorsal stepping index (DSI) is shown in (**b**, Control DSI: 24.41±14.69 vs Dox DSI: 11.75±5.79; *t*=3.383, *df*=7, *p*=.012; individual circles represent individual animal scores at each post-SCI Control and Dox timepoint). Dorsal stepping index accounts for total dorsal steps for both left and right hindlimbs. The total dorsal steps from (b) were separated based on sidedness: right hindlimb (RH) and left hindlimb (LH) (**c**, Control right: 0.568±0.247 vs Dox right: 0.605±0.238, *t*=.554, *df*=7, *p*=.597; Control left: 0.432±0.247 vs Dox left: 0.395±0.238, *t*=.554, *df*=7, *p*=.597; paired t-tests). The right hindlimb showed more dorsal steps overall; however, it maintained that percentage during Dox. No significant differences were seen in sidedness. As with pre-SCI data, circular phase was transformed from a scale of 0-1 to 0.5-1.0 to eliminate lead limb preferences (**d**). Transformed phase values plantar and dorsal steps for hindlimb pair (**e**, Control hindlimbs with dorsal steps: *n*=91/288 [31.60%] vs Dox hindlimbs with dorsal steps: *n*=62/514 [12.06%]; *p*<.001, *z*=7.05) and heterolateral hindlimb-forelimb pair (**h**, Control heterolateral limbs with dorsal steps: *n*=94/288 [32.64%] vs Dox heterolateral limbs with dorsal steps: *n*=160/514 [31.13%]; n.s., *z*=.44, Binomial Proportions tests) are shown. Plantar steps are indicated by grey circles (post-SCI Control) and red circles (post-SCI Dox), while dorsal steps are indicated by teal circles for both post-SCI Control and Dox datasets. The blue boxes indicate values outside of normal variability for the specified uninjured limb pair mean. The percentage of abnormal steps found above normal variability (circles within the blue boxes) is calculated for their respective limb pair (**f,i:** statistics as shown above, B.P test; yellow indicates percentage of steps found within the blue boxes of F and I, respectively). Passes with any left hindlimb dorsal steps were removed and plotted for each of the aforementioned limb pairs (**g,j**: Control hindlimbs without dorsal steps: *n*=33/157 [21.02%] vs Dox hindlimbs without dorsal steps: *n*=39/397 [9.82%]; *p*<.005, *z*=3.13; Control heterolateral limbs without dorsal steps: *n*=59/157 [37.58%] vs Dox heterolateral limbs without dorsal steps: *n*=142/397 [35.77%]; n.s., *z*=0.4; B.P. tests). For the forelimb pair, no significant differences were seen between Control and Dox phase during the first post-injury Dox administration (**k**; PD2 forelimbs: *n*=7/143 [4.90%] vs D2D5 forelimbs: *n*=4/137 [2.92%]; n.s., *z*=.86; PD2 forelimbs: *n*=7/143 [4.90%] vs D2D8 forelimbs: *n*=4/133 [3.00%]; n.s., *z*=.81, B.P. tests). Significance was detected between Control and Dox at the D1D8 time point during the second Dox administration (PD3; D3D8), but no significance was found at the extended Dox time point (D3D20; PD3 forelimbs: *n*=14/145 [9.66%] vs D3D8 forelimbs: *n*=2/135 [1.48%]; *p*<.05, *z*=2.94; PD3 forelimbs: *n*=14/145 [9.66%] vs D3D20 forelimbs: *n*=4/109 [3.67%]; n.s., *z*=1.84, B.P. tests).

We next examined the temporal coordination of the hindlimbs during overground stepping (Fig 5d). Silencing increased the proportion of hindlimb steps that fell within the normal range of left-right phase values, concentrated around 0.5, which reversed when Dox was removed (Fig 5e; PD2, D2D5, D2D8). The improved hindlimb coupling returned when Dox was administered a second time after injury (Fig 5e; PD3, D3D8, D3D20). When compared to pre-SCI controls, the variability in hindlimb-hindlimb and heterolateral hindlimb-forelimb phase values was increased post-SCI with ~33% and ~30% of steps being abnormal, respectively (Fig 5e,f,h,i; grey/yellow). Silencing drastically improved the left-right hindlimb coupling, reducing the number of abnormal steps to ~12% (Fig 5f, pink/yellow). However, the heterolateral hindlimb-forelimb pair saw no significant improvement in coupling (Fig 5h,i; red/pink/yellow).

To control for right-left bias due to behavioral asymmetry, we removed passes that included left hindlimb dorsal steps and re-examined interlimb coordination. Hindlimb coupling was still significantly improved when only plantar steps were considered, despite a reduction in the number of abnormally coordinated steps for both control and Dox time points with this analysis (Fig 5g). The coordination of the heterolateral hindlimb-forelimb pair remained unaltered (Fig 5j). The forelimbs were unaffected and maintained left-right alternation through all conditions (Fig 5k).

Three animals were removed from the data set prior to injury as they showed no perturbations of left-right alternation at any pre-injury Dox time point, presumably due to technical problems with the virus injections (Figure 5 – figure supplement 1a-i). Interestingly, these animals did not show silencing-induced improvements in coordinated stepping post-SCI, suggesting that improvements in hindlimb stepping is an LAPN silencing-induced phenomenon (Figure 5 – figure supplement 1j-l).

### Key features of locomotion are improved during post-SCI LAPN silencing

To explore if silencing spared LAPNs post-SCI similarly improved the speed-dependent spatiotemporal features of hindlimb stepping, we plotted swing time, stance time, and stride distance in relation to speed. These relationships are partially disrupted as a result of SCI, with both plantar and dorsal steps falling outside the normal range (Fig 6a-c). Interestingly, measures that are typically associated with locomotor rhythm (stride time and stride frequency) were also disrupted (Fig 6d,e), but these disruptions were typically associated with a dorsal step, which tends to disrupt interlimb coordination. Silencing spared LAPNs improved these relationships and resulted in a dramatic reduction in the number of abnormal steps, and reduced variability overall, regardless of dorsal or plantar stepping (Fig 6f-j). Average swing (Fig 6k) and stance times (Fig 6l) are improved, with swing time showing the greater improvement. Interestingly, the fundamental characteristic of duty cycle (the ratio of stance time to total stride time) was unchanged after silencing (Fig 6m), suggesting that the improvements in locomotion were strongly focused on interlimb coupling rather than the speed-dependent stance and swing times. Finally, the average stepping speed was unchanged during post-injury silencing (Fig 6n), indicating that increasing speed was not a primary factor in the improved spatiotemporal relationships. Together with the previous findings, these data suggest that silencing LAPNs after SCI is influencing multiple aspects of locomotion lead by interlimb coordination, but also including step cycle spatiotemporal relationships. The result is a greatly improved stepping when spared LAPNs are silenced after SCI.

**Figure 6.**
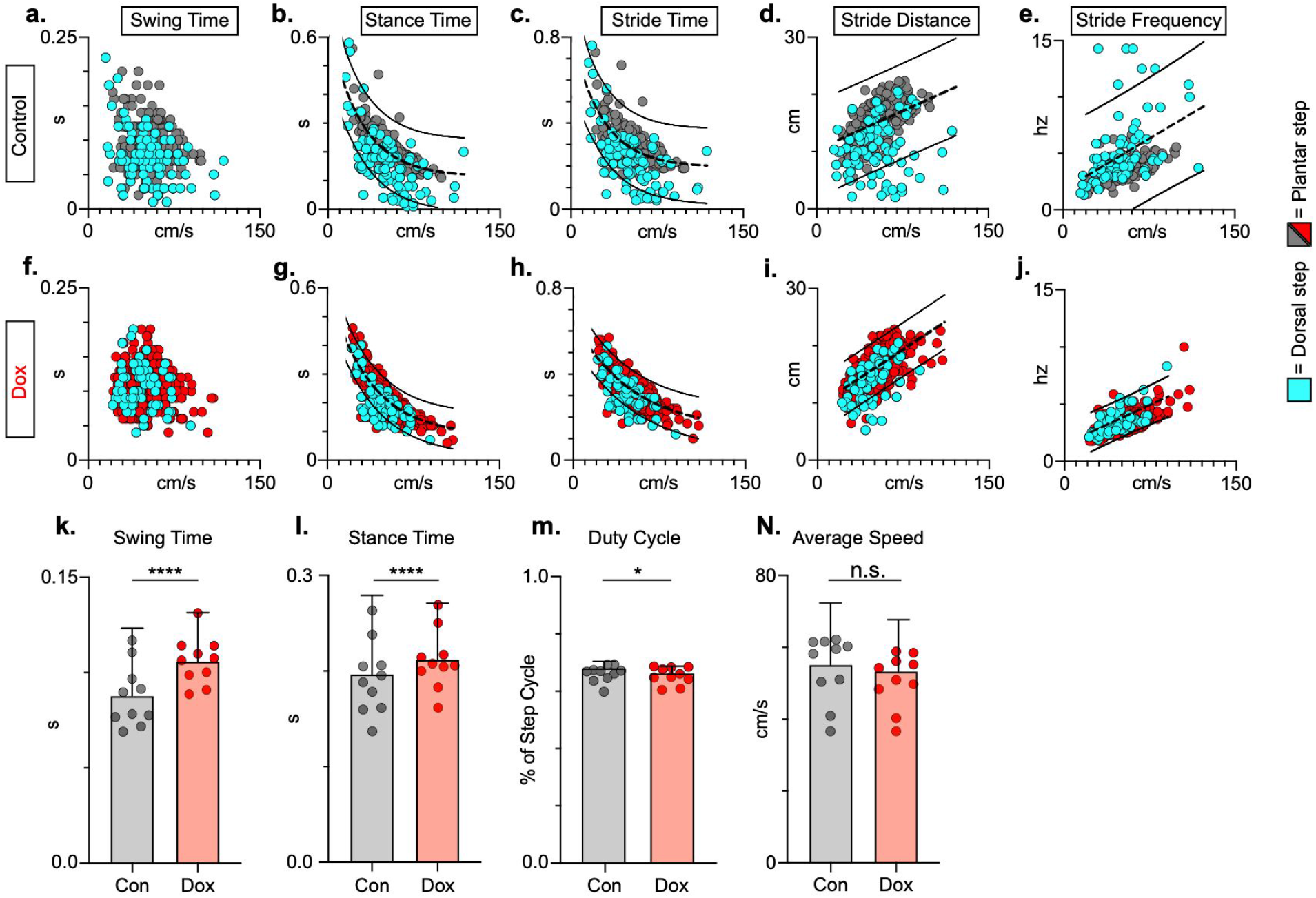
Key features of locomotion are improved following post-SCI LAPN silencing. Relationships between swing time, stance time, stride time, and stride distance are plotted against speed for Control (**a-e**) and Dox (**f-j**) time points. Dorsal steps are indicated with teal circles for both Control and Dox, while plantar steps are indicated with either grey or red circles for Control and Dox, respectively. An exponential decay line of best fit is displayed for stance time and stride time graphs, while a linear line of best fit is displayed for stride distance (dotted line indicates line of best fit; Stance time: Control R^2^=0.433 vs Dox R^2^=0.656; Stride time: Control R^2^=0.351 vs Dox R^2^=0.516; Stride distance: Control R^2^=0.124 vs Dox R^2^=0.367). 95% prediction intervals are shown for lines of best fit as solid lines. Average swing time (**k**, Control swing time: 0.087±0.016 vs Dox swing time: 0.106±0.012, *t*=7.062, *df*=9, *p*<.001, paired t-test) and average stance time (**l**, Control stance time: 0.196±.038 vs Dox stance time: 0.213±0.031, *t*=4.994, *df*=9, *p*=.001, paired t-test) are indicated with circles representing individual animal averages. The average duty cycle (stance time/stride time) (**m**, Control duty cycle: 0.679±0.028 vs Dox duty cycle: 0.663±0.030, *t*=2.678, *df*=9, *p*=.025, paired t-test, likely significant due to tightness of data) and average speed (**n**, Control speed: 55.17±9.20 vs Dox speed: 52.70±7.47, *t*=1.789, *df*=9, *p*=.107, paired t-test) are plotted for Control (grey) and Dox (red) time points with averages indicated by bars. 1 S.D. is shown.

### Hindlimb coordination during swimming post-SCI

Swimming is a bipedal activity where the hindlimbs provide propulsion while the forelimbs steer *(Gruner and Altman, 1980)*. During swimming, the limbs are unloaded and the proprioceptive and cutaneous feedback are different than during stepping. In contrast to our overground findings, silencing LAPNs in the uninjured animal had no effect on left-right hindlimb alternation or any other salient feature of swimming (Fig 7a,b), suggesting that the circuitry responsible for alternation during swimming is lumbar autonomous and does not rely on information carried by the LAPNs. Hindlimb coordination during swimming is strongly disrupted by SCI and surprisingly was modestly improved during post-SCI silencing (Fig 7c). These results suggest that the role of LAPNs in swimming is altered post-SCI, and that the ability of LAPNs to appropriately integrate incoming sensory information from the hindlimbs may play a role in this changed outcome.

**Figure 7.**
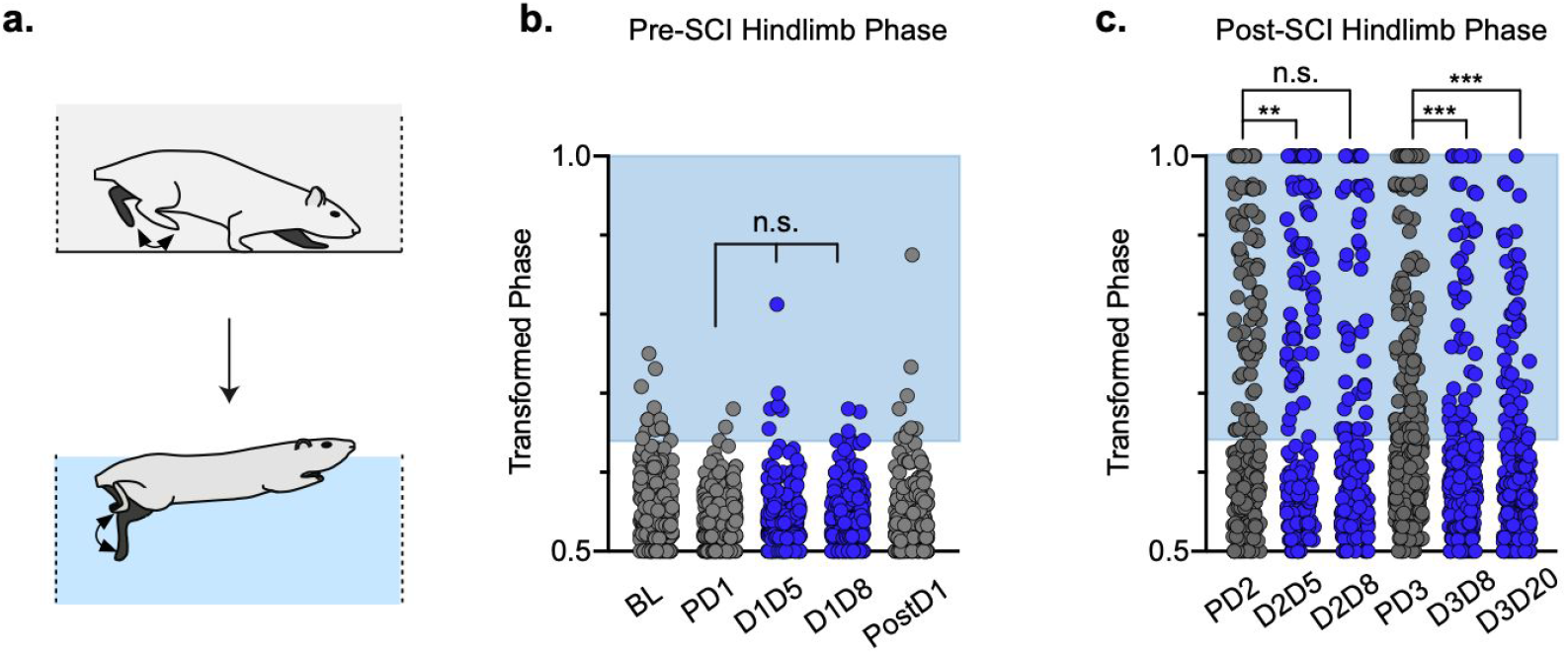
Interlimb coordination is improved during swimming. Similar to hindlimb coordination during overground locomotion, phase can be calculated between the hindlimbs during a swimming task (**a**). Hindlimb alternation was maintained during silencing of LAPNs in the pre-SCI swimming task (**b**, PD1 hindlimbs *n*=4/294 [1.36%] vs D1D5 hindlimbs *n*=8/302 [2.65%]; n.s., *z*=1.13; PD1 hindlimbs *n*=4/294 [1.36%] vs D1D8 hindlimbs *n*=8/278 [2.88%]; n.s., *z*=1.25; grey circles indicate Control swim kicks, blue circles indicate Dox swim kicks). Post-SCI hindlimb coordination was disrupted as a result of injury (**c**, dark grey circles) and was modestly improved as a result of silencing (**c**, blue circles; PD2 hindlimbs *n*=81/188 [43.08%] vs D2D5 hindlimbs *n*=47/166 [28.31%]; *p*=.004, *z*=2.89; PD2 hindlimbs *n*=81/188 [43.08%] vs D2D8 hindlimbs *n*=74/178 [41.57%]; n.s., *z*=0.29; PD3 hindlimbs *n*=119/281 [42.34%] vs D3D8 hindlimbs *n*=59/245 [24.08%]; *p*<.001, *z*=4.42; PD3 hindlimbs *n*=119/281 [42.34%] vs D3D20 hindlimbs *n*=57/233 [24.46%]; *p*<.001, *z*=4.25; Binomial Proportions test). Blue boxes indicate values outside of normal variability for the pre-injury hindlimb pair mean >2 S.D.

## Discussion

Consistent with our previous work, we found that silencing LAPNs in otherwise intact animals disrupted interlimb coordination of both the forelimbs and hindlimbs *(Pocratsky et al., 2020)*. It is intuitive to hypothesize that any spared component of the inter-enlargement pathway represented by the LAPNs should participate in recovered locomotion following an SCI. Unexpectedly, silencing spared LAPNs post-SCI actually improved locomotor function. These results demonstrate modest but meaningful improvements in intralimb coordination when LAPNs are silenced, concomitant with robust improvements in hindlimb interlimb coordination, including paw placement timing and speed-dependent gait indices. Despite these improvements, hindlimb-forelimb coordination improved only slightly during post-SCI silencing.

These findings are counter-intuitive and suggest that the spared LAPNs interfere with hindlimb stepping following thoracic SCI. Given that few LAPNs have local axon collaterals *(Pocratsky et al., 2020)*, these observations suggest that an ascending-descending, inter-enlargement loop involving LAPNs and the descending equivalents, LDPNs, may carry complementary temporal information between the two girdles. The thoracic contusion injury employed significantly reduces the cross-sectional area of white matter at the epicenter, while leaving the outermost rim of the lateral and ventrolateral funiculi intact, precisely where the majority of long-propriospinal axons reside. In addition to frank axon loss, demyelination occurs acutely, and some deficits in remyelination persist chronically, post-SCI *(Totoiu and Keirstead, 2005; Lasiene et al., 2008; Pukos et al., 2019)*, which could lead to slowed and variable action potential conduction velocities. Thus, post-injury, the information carried by the partially spared LAPNs and LDPNs may no longer have the temporal precision necessary to reliably communicate the information needed for interlimb coordination. Rather than aiding in the recovery of stepping, it appears that spared LAPNs hinder the capability of the hindlimb locomotor circuitry below the lesion to function appropriately, contributing to diminished stepping capacity at chronic time points. We speculate that silencing LAPNs may remove excess “noise” within the locomotor system, thereby increasing the ability of intrinsic lumbar circuitry to function independently. Alternatively, maladaptive plasticity may be occurring below the contusion site, leading to detrimental interactions between LAPN input and CPG circuitry.

Another interesting aspect to consider is the loss of influence of LAPNs on forelimb circuitry post-injury, suggesting that their role in providing temporal information helpful for forelimb coordination (alternation) is altered. In our recent study focused on LAPN silencing in uninjured animals, the behavioral phenotype was strongly context-dependent, with disruptions to alternation occurring only during non-exploratory overground locomotion on a walking surface with good grip. Disruptions were insignificant during overground stepping on a slick surface, on a treadmill, or during exploratory-driven stepping *(Pocratsky et al., 2020)*. These observations suggest a dynamic balance between spinal autonomy (where silencing disrupts alternation) and supraspinal oversight (where alternation is maintained). The parsimonious interpretation is that SCI results in increased supraspinal oversight of the cervical circuitry and forelimbs, preventing any disrupted flow of temporal information between the enlargements from disrupting forelimb function. In short, the influence of ascending circuitry post-SCI may be drastically reduced in the presence of increased supraspinal drive. Computational modeling could provide some insight into this conundrum as it would contribute to improved understanding of the drive between the hindlimbs and the forelimbs post-SCI.

Current literature has provided only a basic understanding of LAPN anatomy in the intact spinal cord *(English et al., 1985; Reed et al., 2006; Brockett et al., 2013)*. Therefore, it is unclear whether anatomical and/or transmitter/receptor changes to LAPNs at either the level of the cell bodies in the lumbar cord or at the level of the axon terminals in the cervical cord are contributing to altered behavior after SCI. Further exploration of the anatomical profiles of this population of neurons will be essential to understand improvements seen during LAPN silencing. Another unresolved question is potential silencing-induced plasticity, as improvements in locomotor recovery during silencing were maintained for 20 days. Determining whether synaptic plasticity and/or anatomical changes in LAPNs play a role in improved stepping behaviors during or after silencing will further contribute to our understanding of these perplexing outcomes.

Alternatively, LAPNs may not undergo anatomical remodeling after SCI or silencing and any behavioral changes as a result of silencing could be due to maladaptive plasticity of sensory afferents. The context-specificity of the LAPN silenced phenotype *(Pocratsky et al., 2020)* implies that LAPNs are integrators of hindlimb sensory information. As both proprioceptive *(Beauparlant et al., 2013; Takeoka and Arber, 2019)* and nociceptive *(Krenz and Weaver, 1998)* afferents caudal to the injury sprout post-SCI, newly formed circuits involving these afferents may include aberrant connections making it difficult for the LAPNs to properly integrate the information needed for interlimb coordination. *Ex vivo* studies suggest that the isolated lumbar and cervical enlargements are each capable of producing locomotor-like rhythms and, when in complete spinal preparations, that the lumbar circuitry has a greater influence on cervical circuitry than its reciprocal pathway *(Juvin et al., 2005, 2012; Ballion et al., 2001)*. If LAPNs are receiving aberrant sensory inputs from the hindlimb, removing them from the lumbar and lumbo-cervical circuitry could allow more autonomous correction of the intrinsic patterning within the lumbar cord.

Clinically, epidural stimulation of the lumbosacral spinal cord improves both voluntary motor and locomotor performance in chronic SCI subjects *(Minassian et al., 2007; Harkema et al., 2011; Angeli et al., 2014, 2018)*. In large part, the mechanisms that underlie these outcomes are unknown, but clinically analogous stimulation in rodents was found to activate principally dorsal column and/or primary afferent sensory pathways *(Idlett et al., 2019)*. It is unlikely that clinical epidural stimulation is directly activating lumbar locomotor circuitry or LAPNs, but likely influences lumbar motor circuitry via the more dorsally (posteriorly) located sensory circuitry. This leads us to speculate that the more successful stimulus parameters, chosen via painstaking epidural stimulus mapping *(Mesbah et al., 2017)* may be those that avoid activation of aberrant pathways, including any anomalous sensory input onto LAPNs or lumbar CPG circuitry. It will be critical to consider the presence of some spared pathways, such as this distinct population of LAPNs, as maladaptive to locomotor improvement in humans as it may be contributing to a ceiling effect for human locomotor recovery and to the beneficial effects of epidural and/or transcutaneous spinal cord stimulation.

Collectively, our results demonstrate that some spared axons may be detrimental to locomotor recover after a thoracic contusion, contributing to a “neuroanatomical-functional paradox”, and raising the possibility that neuron- or axon-protective strategies leading to anatomical sparing may not result in the expected benefits *(Fouad et al., 2021)*. Further studies utilizing reverse-engineering strategies to selectively silence/excite identified neuronal populations will be needed to identify which ascending and descending axons are contributory or inhibitory to recovery and the post-injury events leading to these drastic changes in functional role.

## Materials and methods

Experiments were performed in accordance with the Public Health Service Policy on Humane Care and Use of Laboratory Animals, and with the approval of the Institutional Animal Care and Use (IACUC) and the Institutional Biosafety (IBC) Committees at the University of Louisville.

Adult female Sprague-Dawley rats (215-230 g) were housed 2/cage under 12-hour light/dark cycle with *ad libitum* food and water. Each animal served as its own control. Power analysis for gait measures revealed that N=6-10 could detect a true significant difference with power of 85-95%. A total of 16 animals entered the study to account for animal mortality following multiple surgical procedures. N=4/16 animals were excluded for not meeting the *a priori* inclusion criteria based on uninjured behavioral phenotype during LAPN silencing. Of the remaining 12 animals, 2 died after surgery leaving N=10 that were used for the main pre- and post-injury data set.

### Intraspinal injections of viral vectors to doubly infect LAPNs

Virus production, characterization, and injections were done as described previously *(Pocratsky et al., 2017a, 2017b, 2020)*.

### Spinal cord injury

Two weeks after conclusion of uninjured Dox^ON^ assessments, animals were reanesthetized (ketamine:xylazine, 80 mg/kg:4 mg/kg; Henry Schein Animal Health; Akorn Animal Health) and the T9/T10 spine was immobilized using custom-built spine stabilizers *(Hill et al., 2009; Zhang et al., 2008)*. SCI was performed as previously described *(Magnuson et al., 2009)*. Post-injury care during recovery included antibiotics, analgesics, daily manual bladder expression and supplemental fluids as previously described *(Magnuson et al., 2009)*. Animals were allowed to recover for ~6 weeks before any additional pre-DOX assessments.

We utilized a dual viral vector technique to infect LAPNs via their terminals (C6-C7) and cell bodies (L2, Fig 1a). In the presence of doxycycline (Dox), doubly-infected neurons that constitutively express rtTAV16 Tet-On sequence induce expression of enhanced tetanus neurotoxin (eTeNT). At cell terminals, eTeNT prevents synaptic vesicle release, leading to “silenced” neurotransmission. Removing Dox reverses the silencing and restores functional neurotransmission both before and after SCI (Fig 1b) *(Kinoshita et al., 2012)*.

### Experimental timeline

Doxycycline hydrochloride (Dox, 20 mg/ml; Fisher Scientific BP2653-5, Pittsburgh, PA) was dissolved in 3% sucrose and provided *ad libitum* for 8 days pre-injury and for 8-21 days starting 6 weeks post-injury. Dox solution was made fresh, replenished daily, and monitored for consumption. All behavioral assessments were performed during the light cycle portion of the day and concluded several hours before the dark cycle began.

Prior to SCI, behavioral assessments were performed before viral injections (Baseline, “BL”), preceding Dox-induced LAPN silencing (pre-Dox, “PD1”), during Dox (Dox1^ON^D5-D8, “D1D5”, “D1D8”), and 10 days post-Dox (“PostD1”). Animals that met the *a priori* inclusion criteria based on the behavioral phenotype subsequently received SCI. Following SCI, pre-Dox and Dox^ON^ time point assessments were performed twice (Dox2 and Dox3) to assess the reproducibility of the silencing-induced behavioral changes. Data shown are from pre-injury control and Dox timepoints (“pre-SCI”) and post-injury control and Dox timepoints (“post-SCI”). Individual and group comparisons were made for Control vs Dox uninjured and Control vs Dox injured at each time point. Behavioral analyses began on DoxD5 and were repeated on DoxD8. Terminal behavioral assessments occurred on DoxD8 and DoxD21.

### Hindlimb kinematics and intralimb coordination

Acquisition and analysis of hindlimb kinematics were performed as previously described *(Pocratsky et al., 2017a, 2020)*. Briefly, we digitized the movements of the hindlimb using the iliac crest, hip, ankle and toe from two sagittally oriented cameras. Data were exported to a custom Microsoft Excel workbook and hindlimb angles were calculated for each digitized frame. The temporal relationship between the proximal and distal hindlimb angles was calculated for the left and right hindlimbs independently using the peak-to-peak duration of the lead angle during a single step cycle. Stick figures (2D) were generated as previously described *(Pocratsky et al., 2017a, 2020)*.

### Overground gait analyses

Overground gait analyses were performed as previously described *(Pocratsky et al., 2020)*. Dorsal steps, defined by the dorsum of the hindpaw coming into contact with the surface during stance, was considered a step if it maintained contact with the surface during the swing phase of the step cycle. Hindlimb steps were analyzed using the left limb as the reference. As a result, dorsal steps were identified only if they involved the left limb.

Interlimb phase was calculated by dividing the initial contact time of one limb by the stride time (initial contact to initial contact) of the other limb. Phase was represented as circular polar plots to demonstrate interlimb coordination regardless of lead limb and was converted to a linear scale (0.5-1.0) to eliminate any lead limb preferences. When plotted linearly, blue shaded boxes on graphs represent a threshold of >2 standard deviations (S.D.) as calculated from uninjured control average and S.D.

### Open field locomotor assessments

Hindlimb function during overground locomotion was assessed using the BBB Open Field Locomotor Scale, and assessments were performed by trained individuals blinded to experimental time points *(Basso et al., 2002; Caudle et al., 2015)*. Prior to injury, BBB assessments occurred at baseline, pre-Dox, and Dox time points. After SCI, BBB scores were acquired weekly beginning at 7 days post-injury until scores plateaued (~6 weeks post-SCI). BBB testing occurred prior to Dox administration (pre-Dox) and on any days of kinematic testing.

### Stepping coordination indices

Regularity index (RI), central pattern index (CPI), and plantar stepping index (PSI) were used to evaluate gross motor coordination. RI is the number of normal step sequence patterns (NSSP) x4 divided by the number of total cycles. RI excludes any dorsal steps or irregularly patterned steps, and thus is a sensitive measure of stepping quality after SCI *(Hamers et al., 2001)*. CPI is calculated as the number of correctly patterned step cycles divided by the total number of step cycles and includes both dorsal and plantar steps *(Hamers et al, 2001; Koopmans et al., 2005)*. PSI is the number of hindlimb plantar steps divided by the number of forelimb plantar steps. In an uninjured animal, the ratio of hindlimb to forelimb steps is 1:1, or a PSI score of 1.0 *(Magnuson et al., 2009)*. This provides a measure of hindlimb plantar stepping as compared to forelimb stepping after SCI.

### Spatiotemporal indices of stepping

For each individual plantar and dorsal step, temporal and spatial measures were plotted against their instantaneous speed (centimeters/second): swing time (the time the limb is in the air from lift off to initial contact), stance time (the time the limb is in contact with the ground from initial contact to lift off), stride time (stance time + swing time), all in seconds, stride/step frequency (1/stride time), and stride distance (distance traveled per step, centimeters). Average swing, stance and stride times, and stride distances were determined using the individual step values independent of speed. The average speed was calculated from the instantaneous speeds of each step analyzed. Averages were generated for each animal and were plotted with the group average.

### Hindlimb phase during swimming

Swimming assessement *(Pocratsky et al., 2020)* used a maximum of 3 passes with 3-6 complete stroke cycles per pass, per animal in each direction (to the left and to the right). The position of the toe at peak limb extension was digitized for both hindlimbs and the time of peak extension of the nearside hindlimb was divided into the length of time for one complete stroke cycle of the reference opposite hindlimb. Values were transformed as described above and the proportion of strokes with phases >2 S.D. from control mean were compared across time.

### Histological analysis

Animals were euthanized at one of two time points: D3D8 (N=2) or D3D20 (N=11) to determine if viral expression and behavioral changes would persist beyond one week of Dox administration. Histological analysis revealed no differences between D8 and D20 therefore all images are shown from D20 animals.

Animals were overdosed with sodium pentobarbital, followed by pneumothorax and transcardial perfusion with 0.1 M phosphate-buffered saline (PBS) (pH 7.4) followed by 4% paraformaldehyde in PBS. Spinal cords were dissected, post-fixed for 1.5 hours, and transferred to 30% sucrose for > 4 days at 4°C. Spinal segments C5-C8, T8-T12, and T13-L3/L4 were dissected, embedded in tissue freezing medium, and stored at −20°C until they were cryosectioned at 30 *μ*m.

EGFP.eTeNT expression in the cervical spinal cord was confirmed immunohistochemically. Sections were incubated at 37°C for 30 minutes, then were rehydrated in PBS for 10 minutes (pH 7.4, room temperature) followed by a 60 minute incubation in a blocking solution made of nine parts milk solution (Bovine Serum Albumin (BSA), 0.75g of powdered skim milk, and 14.25ml 0.1% of PBS with Tween 20 (PBST)) and one part 10% Normal Donkey Serum (NDS)). This was followed by another 10-minute wash in PBS and overnight incubation at 4°C in milk solution containing primary antibodies. The primary antibody milk solution contained a combination of rabbit anti-GFP (to enhance eTeNT.EGFP) and either guinea pig anti-synaptophysin (presynaptic marker), guinea pig anti-vesicular glutamate transporter 1 (VGlut1, sensory afferents), guinea pig anti-vesicular glutamate transporter 2 (VGlut2, excitatory synapses), or goat anti-vesicular GABA transporter (VGAT, inhibitory synapses). For information on primary antibodies, see Key Resources Table.

On Day 2, tissue sections were washed several times at room temperature, alternating between PBS and 0.1% PBST, followed by a 1-hour incubation in milk solution containing the following secondary antibodies in a dark room: donkey anti-rabbit Alexa-Fluor Plus 488 (1:500), donkey anti-guinea pig Alexa-Fluor 594 (1:200), and donkey anti-goat Alexa-Fluor 594 (1:200) (Key Resources Table). Tissue sections were washed with PBS for 10 minutes and then incubated at room temperature with fluorescent Nissl (NeuroTrace 640/660 Deep Red, ThermoFisher N21483, dilution of 1:100) in PBS for 1 hr to stain neuronal cell bodies. Tissue was then washed with PBS for 2 hrs and coverslipped using Fluoromount (Southern Biotechnology Associates, Inc.; Birmingham, AL, USA). The above procedure was repeated on lumbar spinal cord sections. Isotype matched IgG with identical protein concentration was used as a negative control (donkey anti-rabbit IgG; Jackson ImmunoResearch #711-005-152).

Fluorescent images were captured on an Olympus Fluoview 1000 confocal microscope. Tissue sections were viewed with an oil immersion 100x objective using 488, 543, and 647 lasers (Olympus; Center Valley, PA, USA). Z-stacks ranged from 55-64 slices at 0.45 *μ*m optical steps. Neurons within the intermediate grey matter were imaged for both cervical and lumbar sections. Images were analyzed using Amira software as described *(Pocratsky et al., 2017a, 2020)*.

Spared white matter at the injury epicenter was assessed as described previously using sections stained with iron eriochrome cyanine and alkali differentiation (EC) *(Magnuson et al., 2005)*. Briefly, slides were allowed to warm at room temperature for 60 minutes before being placed in a hydration gradient consisting of xylenes, ethanol, and distilled water. EC stain was applied to the slides for 10 minutes, followed by two short applications of distilled water to remove excess stain (10-15 seconds). Slides were placed in differentiating solution for 30 seconds before air drying overnight. On Day 2, slides were placed in xylenes solution for 10 minutes and were coverslipped with Permount. Sections were imaged using a Nikon Eclipse E400 light microscope at 10x magnification. Spot Software (v.5.1) was used to format images.

### Statistical analyses

Statistical analyses were performed using the SPSS v22 software package from IBM. Additional references for parametric and non-parametric testing were used *(Hays, 1981; Siegel and Castellan, 1988; Ott, 1977)*. Differences between groups were deemed statistically significant at p≤.05. Two-tail p-values are reported.

The Binomial Proportion Test was used to detect significant differences in the proportion of coordination values beyond control threshold (>2 S.D.) for the raw and transformed values of interlimb coordination of various limb pairs prior to and post-SCI. It was also used to determine statistical significance for interlimb phase, raw BBB score differences, intralimb phase, dorsal steps as a percentage of total steps, and percentage of categorically organized steps (anti-phase, out of phase, in phase). Statistical outliers were excluded when appropriate.

Circular statistics were performed on the stepping interlimb coordination datasets *(Pocratsky et al., 2017a; Zar, 1974)*. We primarily used the non-parametric two-sample U^2^ test based on a previously described rationale *(Pocratsky et al., 2017a, 2020)*. The null hypothesis tested was whether two time points have the same expression of coupling pattern (right-left phase), i.e. phase values are concentrated around 0.5 or they are distributed throughout the possible range.

Regression analyses to compare the slopes for the lines of best fit were performed on the speed versus spatiotemporal gait indices datasets, including speed vs. swing, stance and stride times, and stride distance for both hindlimb and forelimb pairs before and after SCI. For regression analyses post-SCI, plantar and dorsal steps were included in the analysis and dorsal steps are shown in teal. 95% prediction intervals are indicated on the graphs by solid lines. Dorsal steps were never seen for the forelimbs, thus all forelimb regression analyses were performed on plantar steps only.

Mixed model analysis of variance (ANOVA) followed by Bonferroni *post hoc* t-tests (where appropriate) were used to detect a significant difference in the BBB scores based on sidedness and condition (i.e. Control vs Dox; data not shown,). No significant differences were observed between sidedness and within condition. Repeated measures ANOVA were used to ensure that the average number of steps analyzed per animal during uninjured and injured time points. No differences were detected (for all comparisons, p<.05).

Paired t-tests were used to detect significant differences in proximal and distal angle excursion for intralimb coordination, average gross stepping measures including RI, CPI, PSI, and BBB scores, average intralimb phase, percentage of dorsal step sidedness, average swing time, average stance time, average duty cycle, and overall average speed at Control and Dox pooled time points, respectively.

## Abbreviations

BBB: Basso, Beattie, and Bresnahan Locomotor Scale
BSA: bovine serum albumin
CPGs: central pattern generators
CPI: central pattern index
Dox: Doxycycline
DSI: dorsal stepping index
EC: eriochrome cyanine
eTeNT: enhanced tetanus neurotoxin
IACUC: Institutional Animal Care and Use Committee
IBC: Institutional Biosafety Committee
LAPNs: long ascending propriospinal neurons
LDPNs: long descending propriospinal neurons
NDS: normal donkey serum
NSSP: normal step sequence patterns
PSI: plantar stepping index
RI: regularity index
SCI: spinal cord injury
VLF: ventrolateral funiculus

## Acknowledgements

The authors thank Dr. Tadashi Isa and Dr. Akiya Watakabe for generously providing the viral vector plasmids, Russell M. Howard for assistance in viral vector production, Jason E. Beare for assistance in confocal imaging and image processing, and Christine Yarberry, Kariena Andres, and Alice Shum Siu for their surgical support and assistance with animal care.

---

**Figure 1 – figure supplement 1.**
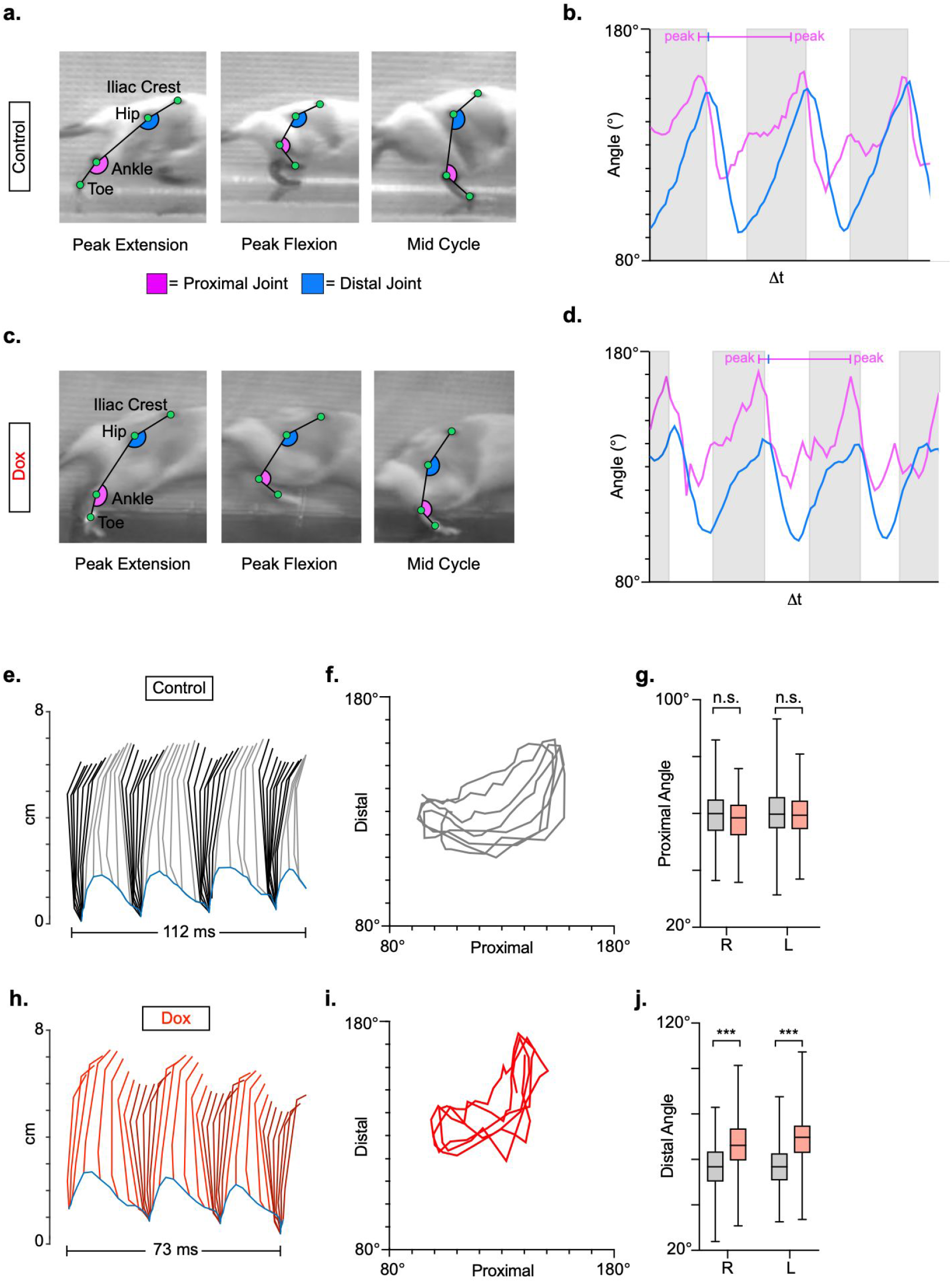
LAPNs do not contribute to intralimb locomotor coordination of uninjured animals. Representative images of intralimb joint angles of the proximal (iliac crest–hip–ankle) and distal (hip–ankle–toe) joints during Control (**a**) and Dox (**C**) timepoints. Example images are taken from 3 moments within a step cycle: peak extension, peak flexion, and mid stance. We examined the peak excursion of each angle at Control and Dox timepoints using excursion traces of the proximal and distal joint angles, indicated by the blue and purple lines, respectively (**b,d**). Grey boxes indicate stance portions of the step cycle. Two-dimensional stick figures of hindlimb stepping (**e,h**) and angle-angle plots (**f,i**; sampling rate = 100 frames/sec) from the same passes shown in (**a,c**). Toe height throughout the pass is indicated as a blue trace on the bottom of the two-dimensional stick figures. The average excursions of the proximal angles (**g**, Control right: 58.79±4.94 vs Dox right: 56.56±3.12, *t*=2.08, *df*=11, *p*=.062; Control left: 60.40±4.04 vs Dox left: 58.65±4.23, *t*=2.17, *df*=11, *p*=.053; paired t-tests) and distal angles (**j**, Control right: 55.84±4.93 vs Dox right: 65.57±8.46, *t*=4.55, *df*=11, *p*=.001; Control left: 56.19±4.24 vs Dox left: 66.64±7.16, *t*=6.63, *df*=11, *p*<.001; paired t-tests; center bars indicate mean with extending bars indicating range of data) for control and silenced timepoints are shown.

**Figure 5 – figure supplement 1.**
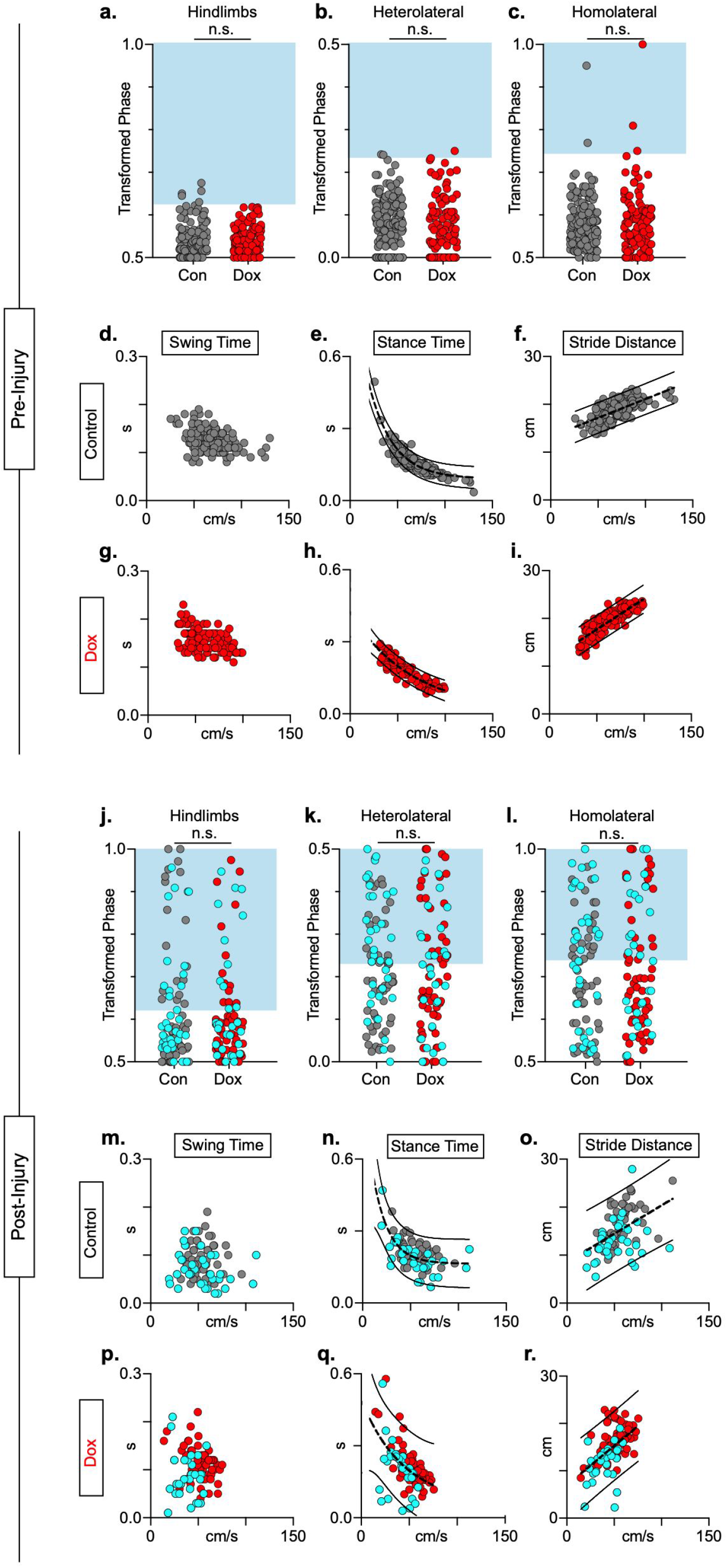
Animals excluded based on lack of behavioral outcomes pre-SCI show no improvements in hindlimb coupling post-SCI. N=3 animals were excluded from post-injury group data based on lack of behavioral phenotype at any time point during pre-injury silencing. No disruptions were seen in hindlimb-hindlimb (**a**, Control hindlimbs: *n*=5/160 [3.13%] vs Dox hindlimbs: *n*=0/110 [0%]; n.s., *z*=1.87), heterolateral hindlimb-forelimb (**b**: Control heterolateral: n=3/160 [1.88%] vs Dox heterolateral: n=3/110 [2.72%]; n.s., *z*=.45), homolateral hindlimb-forelimb (**c**; Control homolateral: *n*=2/160 [1.25%] vs Dox homolateral: *n*=3/110 [2.72%]; n.s., *z*=.83), or forelimb-forelimb (data not shown) phase values during silencing. Blue boxes indicate values outside of normal variability for the uninjured hindlimb pair mean >2 S.D. No changes were seen in relationships between swing time (**d,g**), stance time (**e,h**), or stride distance (**f,i**) prior to injury. Stride time and stride frequency have been omitted; however, no significant differences were observed. After SCI, improvements were not seen in hindlimb-hindlimb phase (**j**: Control hindlimbs: *n*=30/87 [34.48%] vs Dox hindlimbs: *n*=24/87 [27.57%]; n.s., *z*=.99) and similar disruptions in heterolateral hindlimb-forelimb (**k**: Control heterolateral: *n*=43/87 [49.42%] vs Dox heterolateral: *n*=38/87 [43.68%]; n.s., *z*=.76) and homolateral hindlimb-forelimb (**l**, Control homolateral: *n*=41/87 [47.12%] vs Dox homolateral: *n*=33/87 [37.93%]; n.s., *z*=1.23) pairs were unchanged during silencing. Lack of improvement was seen in spatiotemporal measures during silencing (**m-o**, **p**-**r**). Control plantar steps = grey circles, Dox plantar steps = red circles. All dorsal steps are shown as teal circles.

**Video 1**. Injured Control example “bad” pass from side camera view and ventral view (1x speed and 0.25x speed)

**Video 2.** Injured Dox example “good” pass from side camera view and ventral view (1x speed and 0.25x speed)

